# Horizontal gene transfer barrier shapes the evolution of prokaryotic pangenomes

**DOI:** 10.1101/2020.04.14.041392

**Authors:** Itamar Sela, Yuri I. Wolf, Eugene V. Koonin

## Abstract

The genomes of bacteria and archaea evolve by extensive loss and gain of genes which, for any group of related prokaryotic genomes, result in the formation of a pangenome with the universal, asymmetrical U-shaped distribution of gene commonality. To elucidate the evolutionary factors that define the specific shape of this distribution, we investigate the fit of simple models of genome evolution to the empirically observed gene commonality distributions and genomes intersections for 33 groups of closely related bacterial genomes. The combined analysis of genome intersections and gene commonality shows that at least one of the two simplifying assumptions that are usually adopted for modeling the evolution of the U-shaped distribution, those of infinitely many genes and constant genome size, is invalid. The violation of both these assumptions stems from the horizontal gene transfer barrier, *i.e*. the cost of accommodation of foreign genes by prokaryotes.

## Introduction

With the accumulation of complete prokaryotic genomes, it has become evident that even closely related prokaryotes can substantially differ in their gene repertoires [1, 2]. Accordingly, for a collection of genomes, often, collectively construed as a species, it is natural to consider the pangenome, which is defined as the entire non-redundant gene repertoire spanned by the constituent genomes [3]. Reconstruction of the evolutionary dynamics of microbial pangenomes is essential for understanding the evolution of the traits of microbes including ecology, pathogenicity and resistance.

The pangenome consists of genes of widely different abundances. Roughly, the genes in a pangenome can be divided into three classes according to their abundance: 1) the core, which is the collection of genes that are present in (nearly) all genomes, 2) the moderately conserved ‘shell’, and 3) the ‘cloud’ of rare and unique genes [4]. To analyze genes abundances and their evolution quantitatively, it is convenient to analyze the distribution of gene commonality [5-9]. Gene commonality, *g_k_*, is defined for a collection of *N* genomes as the number of genes that are present in exactly *k* genomes, where *k* = 1,2,…, *N*. The distribution of gene commonality is typically U-shaped, with different degrees of asymmetry (Fig. 1a), where the right peak corresponds to the conserved core, the shallow region in the middle to the moderately conserved shell, and the left peak to the cloud of rare genes [4]. In this representation, the pangenome is given by the sum of all points.

**Figure 1.**
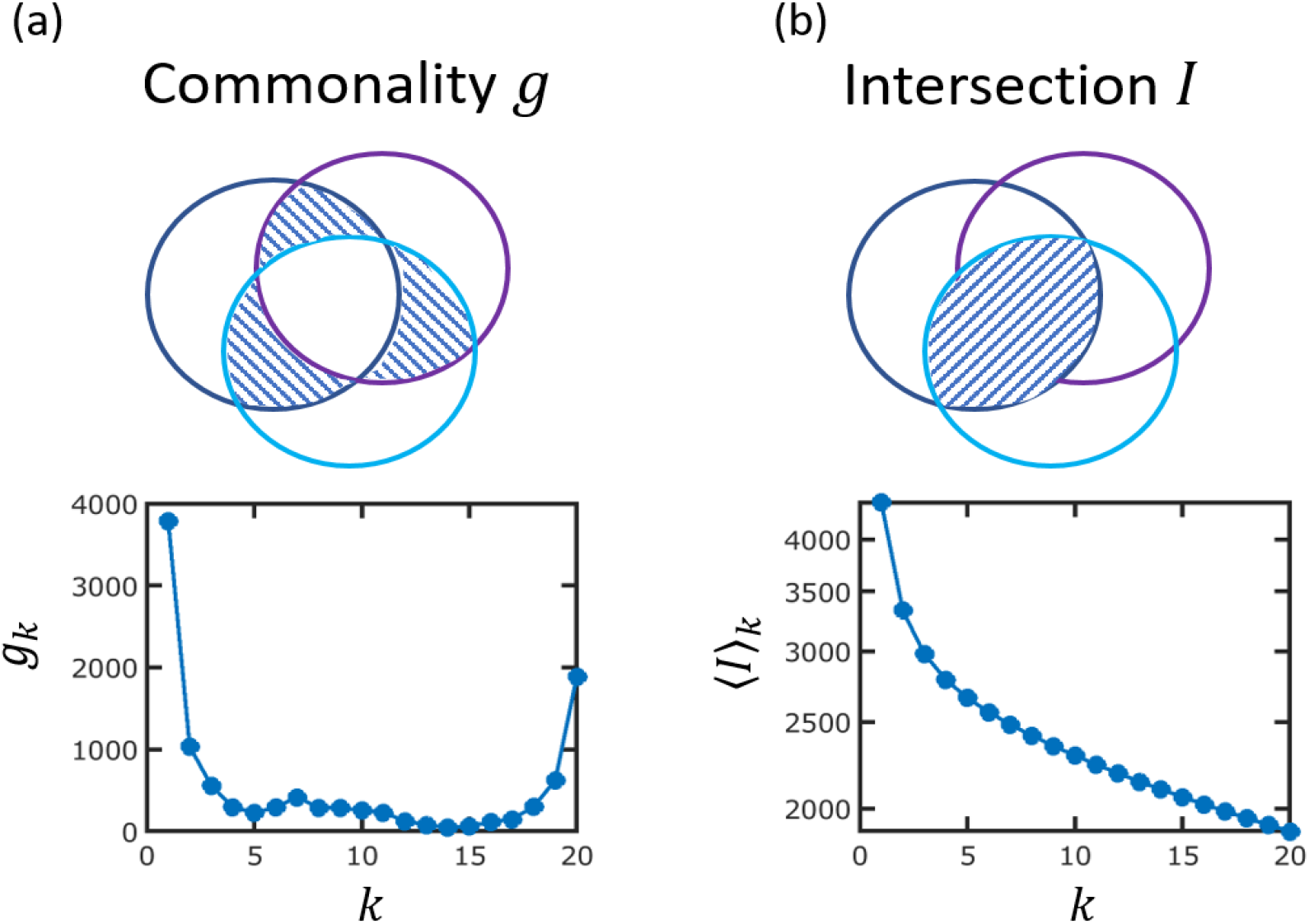
Gene commonality and genome intersection. The Venn diagrams illustrate the overlap of gene contents between genomes. The shaded areas demonstrate the difference between genes commonality (panel **a**) and genomes intersection (panel **b**). Note that. whereas *g_2_* (intersection between two genomes) is a single number, for three genomes, there are three pairwise intersections, and only one of the three possibilities is indicated by the shaded area. In panel **a**, the common gene core of the three genomes is shown by the white are in the middle.

Prokaryotic genome evolution involves extensive gene loss and horizontal gene transfer (HGT) [10-14], which result in the formation of the characteristic structures of the pangenome and the core-genome [15, 16]. Accordingly, the evolutionary models that account for the shaping of the gene commonality U-shape distribution incorporate genes gain and loss rates. In a pioneering work of Baumdicker *et al*. [5], genes commonality was calculated under the assumption that genomes evolve around the equilibrium point where gain and loss rates are equal, such that the genome size is roughly constant. Inspired by the infinitely many alleles model of Kimura and Crow [17], the genes abundance was inferred under the assumption of the infinitely many genes (IMG) model. Under IMG, it is assumed that genes are acquired exclusively from an external, infinite gene pool, and every gene acquisition event introduces a new gene into the genetic repertoire. The IMG assumption was further implemented in different evolutionary models to study the formation of the U-shape distribution in clusters of closely related bacterial genomes. It has been shown that at least two turnover rates of genes are required to accurately account for both the size of the core genome and the number of singleton genes [7]. Genes commonality was also studied in a general context of shared components in complex systems [8]. By considering complex systems of widely different nature and origins, including bacterial genomes, language texts, and Lego kits, it has been shown that the distribution of shared components has universal characteristics. In this work, genomes were regarded as random samples of genes, and the exact phyletic relations of the genomes were not incorporated into the analysis. However, more recently, it has been demonstrated that phyletic patterns could be inferred from the gene commonality distribution [18].

In previous studies [5-7], model parameters, in particular, gene turnover rates, were inferred from the gene commonality distribution. This approach requires explicit assumptions regarding the gene gain and loss rates. For example, the IMG model assumes that the gain rate is constant whereas the loss rate is proportional to the genome size [5, 6]. More importantly, fitting the model directly to gene commonality distribution can result in inaccurate inference of turnover rates. Specifically, as we show in this work, the inference of the turnover rates from the gene commonality distribution will result in under-estimation of the turnover rates due to the breakdown of the model assumptions.

Alternatively, gene turnover rates can be extracted directly from genome intersection *I_k_*, which is defined as the number of genes that are common to (at least) *k* genomes. Genome intersection is thus distinct from gene commonality *g_k_*, which is given by the number of genes that are present in exactly *k* genomes (Fig. 1). We have recently shown that genome intersection decays exponentially with the evolutionary distance [19, 20]. These findings are consistent with multiple pairwise comparisons of genomes in archaea [21], bacteria [22] and bacteriophages [23].

Here, we analyze the divergence of prokaryotic genomes within the theoretical framework we developed previously [19, 24] and extract gene commonality from genomes intersection using the inclusion-exclusion principle [25]. This allows us to obtain the observed U-shape distribution without assuming any specific functional form for the gain and loss rates. However, similar to the IMG model, it is assumed that gain and loss rates are equal, and that genes are gained from an external infinite gene pool. Analysis of gene commonality and genome intersections in 33 groups of closely related prokaryotes indicates a greater number of gene losses compared to gene gains rate. This observation implies breakdown of one or both of the model assumptions: either the actual gene gain rate is smaller than the loss rate, such that genomes shrink with time, or/and genes that are already present in the pangenome are often regained from the external pool that, in such a case, cannot be considered effectively infinite. Both these deviations from the model assumptions appear to be manifestations of the HGT barrier [26], that is, the cost of integrating new genes into the functional networks that already exist in the recipient organism.

## Results

### Extraction of genes commonality from genomes intersections

Prokaryotic genome evolution is dominated by gene loss and HGT [10-14]. It is therefore natural to model genome evolution as a stochastic process where genes are gained and lost at random, with rates *P*^+^ and *P*^−^, respectively [24]. Within this modeling framework, the intersection of *k* genomes, *I_k_*, decays exponentially with the total evolutionary distance *D_k_* (see Methods for details). The exponential decay constant, *λ*, is given by the per-gene loss rate *λ~ P*^−^/*x*, where *x* is the number of genes [19]. The exponential decay of genome intersections is obtained under two assumptions. First, it is assumed that gain and loss rates are closely similar, such that the number of genes is roughly constant. Second, it is assumed that genes are acquired from an infinite gene pool, such that each gain expands the pangenome by one new gene (hereafter Infinite Gene Pool and Constant Genome Size, or IGP-CGS assumptions).

Next, we have to take into account the evolutionary relationships among the intersecting genomes in a given cluster and calculate the mean intersection 〈*I*〉 for each *k*. Evidently, in a cluster of *N* genomes, a subset of *k* genomes can be chosen in multiple ways (see Methods). The evolutionary relationships among the genomes are described by a phylogenetic tree, and for the calculation of 〈*I*〉*_k_*, the mean evolutionary distance for each *k* is weighted according to the tree topology (see Methods for explicit formulation). Using the inclusion-exclusion principle [25], it is possible to extract the gene commonality *g_k_* from the mean genome intersectionwhere 〈*I*〉_*k*_

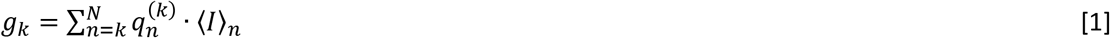

where

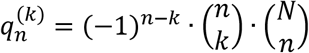

It should be noted that this relation is exact and does not rely on any approximation or assumption involved in the derivation of *I_k_*. This relation is not only a formal result but also provides an intuition with respect to the formation of the U-shape distribution.

Moreover, the combined analysis of genome intersections and gene commonality, and in particular, the relation encapsulated in Eq. 1, carries signatures of the evolutionary dynamics. For the core genome *k* = *N*, and the sum of Eq. 1 contains only one term *g_N_* = 〈*I*〉_*N*_. The core is given by the intersection that is associated with the longest evolutionary time, and its size determines the number of gene losses. For *k* < *N*, gene commonality is calculated backwards from the core genome, in an alternating sign series of *N* − *k* + 1 terms (see Eq. 1). The number of singletons *g_±_* is given by a sum of *N* terms, with all intersections for *k* > 1 subtracted or added to the term *N* · 〈*I*〉_1_, where 〈*I*〉_1_ is the mean genome size. The number of singletons is therefore strongly sensitive to the genome size which, in the course of evolution, is maintained by gene gain. Under the CGS assumption, the number of gain events is equal to the number of loss events. Further assuming an infinite gene pool implies that all gained genes are initially singletons and contribute to *g*_±_. This means that the size of 〈*I*〉_1_ remains constant whereas all other 〈*I*〉_*k*_ with *k* > 1 are shrinking due to gene loss. As we show in the following section, extraction of gene commonality from genomes intersections using the IGP-CGS model results in overestimation of the number of singletons although the IGP-CGS model yields an excellent fit to the mean genomes intersections (*R*^2^ > 0.996). This discrepancy implies violation of the IGP-CGS assumptions. When the IGP assumption does not hold, genes can be regained from the finite external pool resulting in an increase of 〈*I*〉_*k*_ with *k* > 1 and a smaller number of singletons. Violation of the CGS assumption results in a decrease of the genome size, i.e. a lower 〈*I*〉_1_ value, and a a smaller number of singletons as well), even for the large number of gene losses that is implied by the size of the core genome.

### Analysis of simulated genome datasets

To demonstrate the calculation stages, we perform the analysis for a simulated dataset (Fig. 2). Specifically, we aim to demonstrate the stages of gene commonality extraction from genomes intersections. To simulate the evolution of genome content, we represent a genome as a collection of genes. Starting from the root and propagating along the tree branches, genes are lost and gained stochastically, according to the set gain and loss rates (for the complete description of the simulation scheme, see Methods). The simulations were performed for 20 genomes using the tree of the *E. coli* cluster (Fig. 2a), and under the IGP-CGS assumptions.

**Figure 2.**
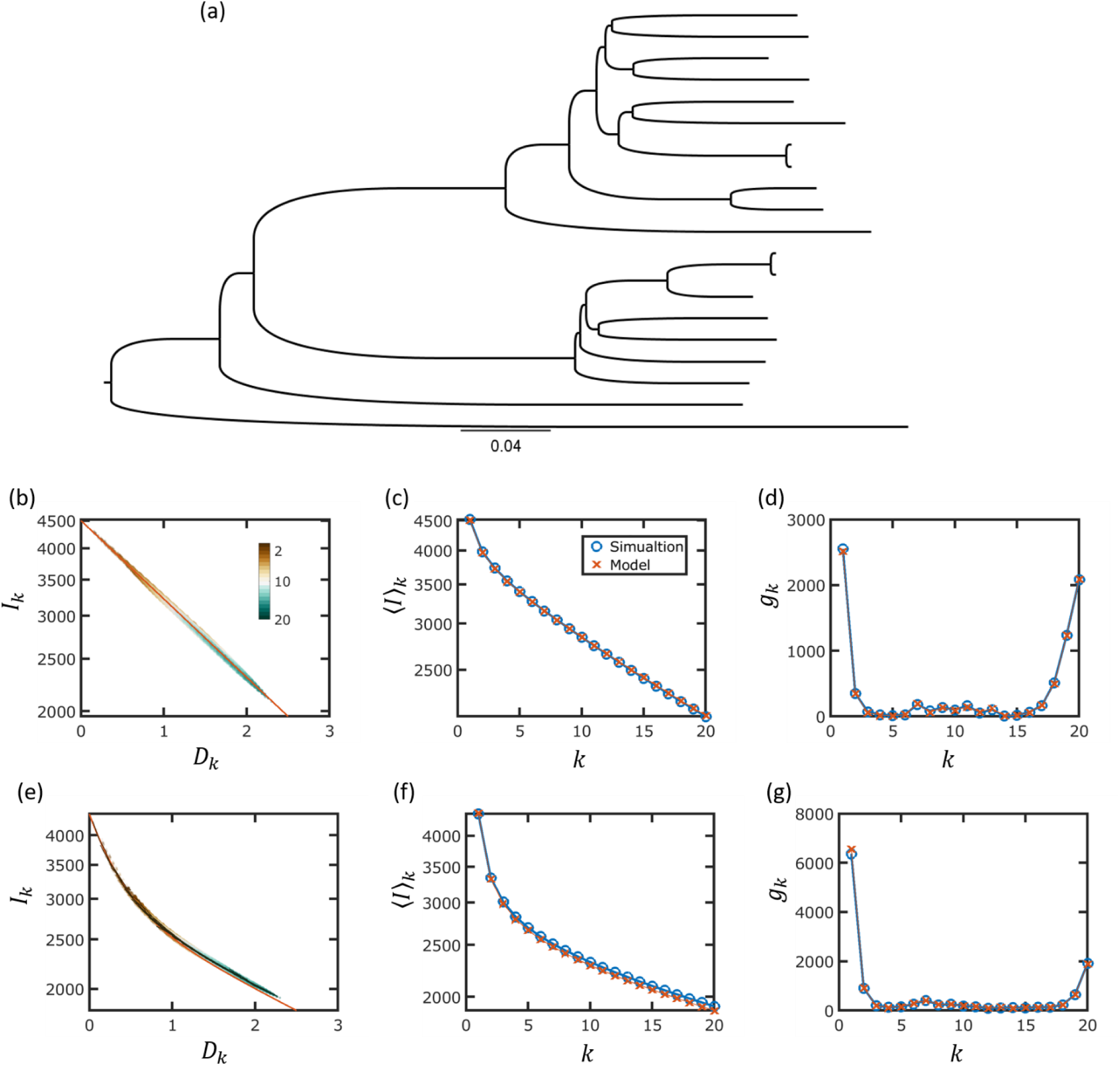
Estimation of genome intersection and gene commonality for a simulated genome dataset. **a)** The phylogenetic tree used for generating the simulated dataset. b) Genomes intersections for a single turnover rate. Numbers of intersecting genomes are shown in different colors, as indicated by the colorbar. The parameters used for the simulations are *x* = 4500 and *P*^+^ = *P^−^* = 1.5 × 10^3^. Model prediction is obtained by substituting *x* and *P^−^* into Eq. 3. Model prediction is indicated by a red line. **c)** Mean genome intersections are calculated for the tree using Eq. 5. The simulated dataset is compared to model prediction (see legend). **d)** Genes commonality for the simulated dataset and model prediction. The model prediction for genes commonality is extracted from the mean genomes intersections of panel **b**, using Eq. 1. Panels **e-g:** simulations using two turnover rates. Model parameters are identical to the values that were inferred for *E. coli* (see Table S1). Panels content is analog to panels **b-d**. Model prediction for *I_k_* was obtained using Eq. 4.

Figs. 2b-d show the results of a single simulation using one gene turnover rate. The genome intersections and gene commonality values are extracted directly from the simulated dataset. For comparison, the analytical results are shown as well. In Fig. 2b, intersections are shown as a function of the evolutionary distance, as computed from the tree that was used for the simulation. For a single gene turnover rate, the intersections decay exponentially and therefore appear as straight lines in the semilog plot of Fig. 2b. Next, the mean intersections 〈*I*〉_*k*_ are shown for all *k*’s in Fig. 2c. For the simulated dataset, 〈*I*〉_*k*_ is calculated for each *k* by taking the mean of all intersections *I_k_*. The model prediction for 〈*I*〉_*k*_ is obtained by averaging the exponential decay expression for *I_k_*, with weights that correspond to the tree topology (see Methods). Finally, the distribution of gene commonality *g_k_* is shown in Fig. 2d, where model prediction for *g_k_* is extracted from the mean genome intersections 〈*I*〉_*k*_ using Eq. 1.

Notably, the U-shape distribution of gene commonality is observed for a single decay constant (Fig. 2d), where all genes evolve at the same rate such that no selection is involved in the pangenome evolution [9]. In the absence of selection, the interpretation of the pangenome evolutionary trajectory is as follows. Initially, at the root of the tree, all genomes are identical, and therefore, all genes belong to the core. In the course of evolution, any gene loss event reduces the size of the core. In other words, due to the loss events, genes move from *k* = *n* to *k* = *n* − 1 occurrence, such that the core can be regarded as a source that diffuses to the left by gene loss. For short enough evolutionary times, a substantial fraction of the original core genome is retained, forming the observed right peak in Fig. 2d. On average, every time a gene is lost, a gene is gained. Due to the infinite gene pool assumption, each time a gene is gained, it is initially a singleton and contributes to *k* = 1 abundance. The gained genes form the left peak that consists of singletons and can be regarded as a source that diffuses to the right from *k* = *n* to *k* = *n* + 1 occurrence through strain divergence and speciation.

To further explore the generality and validity of our modeling scheme, we repeated the entire procedure for a simulation with two gene-turnover rates (Figs. 2e,f,g)). As shown in the next section, two turnover rates were required to fit the model prediction for genome intersections, corresponding to two classes of genes, fast evolving and slow evolving ones. We therefore simulated the evolution of a genome that contains two classes of genes. In this case, the intersections (Fig. 2e), are given by a sum of two exponents (see Methods. The simulation was performed using realistic model parameters that were inferred from the analysis of the *E. coli* genome cluster of (see Table S1).

### Fitting the model to the genomic data

After establishing the analysis strategy using simulated datasets, we applied the same calculation scheme to the genomic data. Specifically, we inferred gene turnover rates from genome intersections and compared the model predictions with the empirical gene commonality distributions. The analysis was performed with an extensive dataset that consisted of 33 clusters of closely related prokaryotic genomes (see Methods).

In accord with the notion of core and accessory genes [16, 27], analysis of the genomic data shows that at least two turnover rates are required to fit the data (Fig. 3 and Figs. S1-S6), indicating that genomes can be roughly divided into slow (core) and fast (accessory) evolving genes. The turnover rate of the fast genes is roughly 10 times greater than that of the slow genes (see Fig. 4 and Table S1). In terms of genome evolution, the fast or slow turnover rates reflect the different average selective effects associated with gene deletion: losing a slow-turnover gene is associated with greater fitness cost than losing a fast-turnover gene [20]. The fraction of fast evolving genes varies from as low as 7% up to 40%, apparently, reflecting substantial differences in the pangenome dynamics among prokaryotes (Fig. 4).

**Figure 3.**
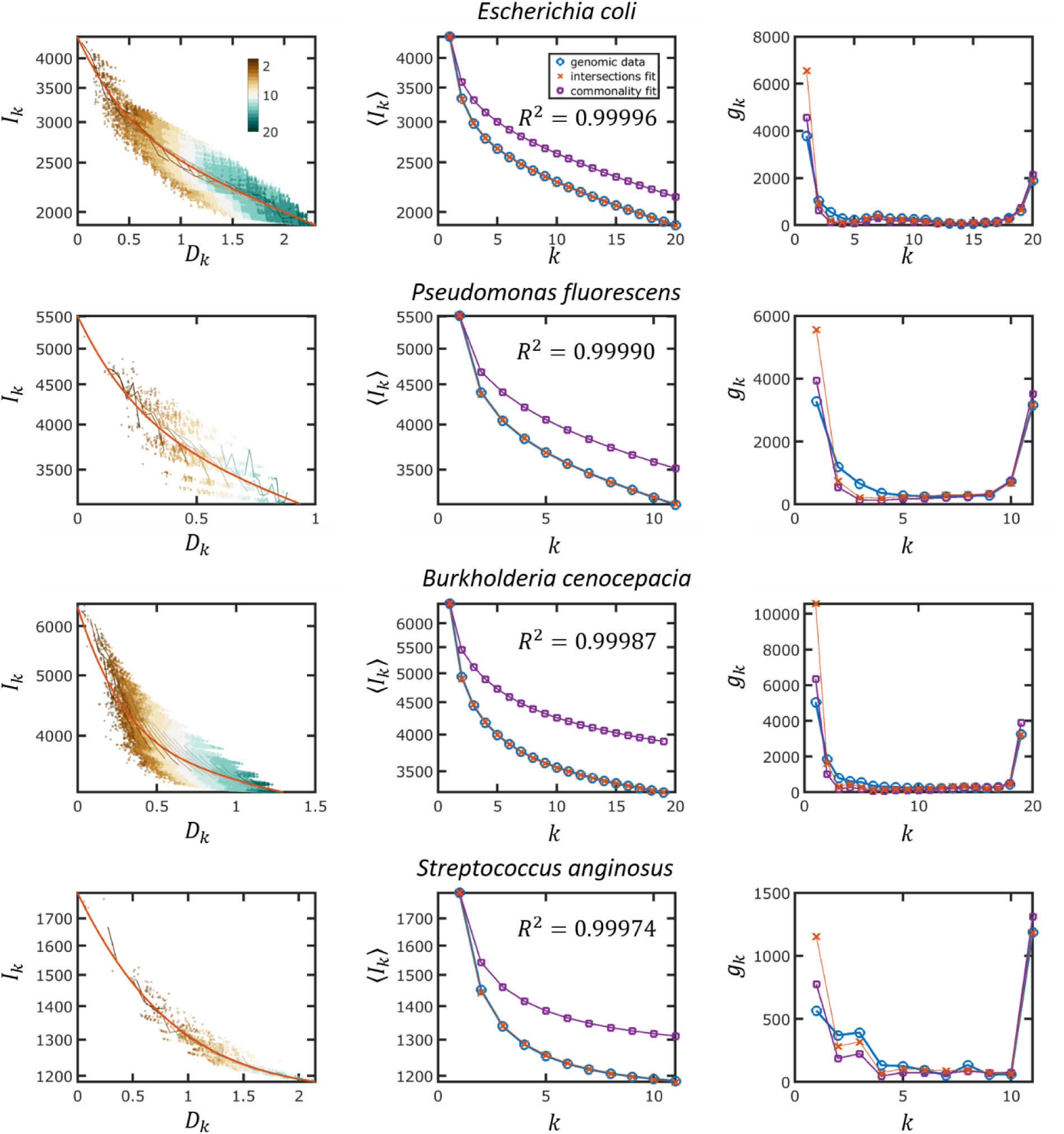
Empirical data and model fits of genome intersection and gene commonality for four genome clusters. The most common species in the cluster is indicated for each cluster. The panels on the left show genome Intersections, the panels in the middle show mean genomes intersections, and the panels on the right show gene commonality (U-shaped distributions). The genomic data are shown by blue circles, and the model fits are shown by red x’s (genome intersection fit) or purple squares (gene commonality fit). The goodness of fit (*R^2^*)is indicated.

**Figure 4.**
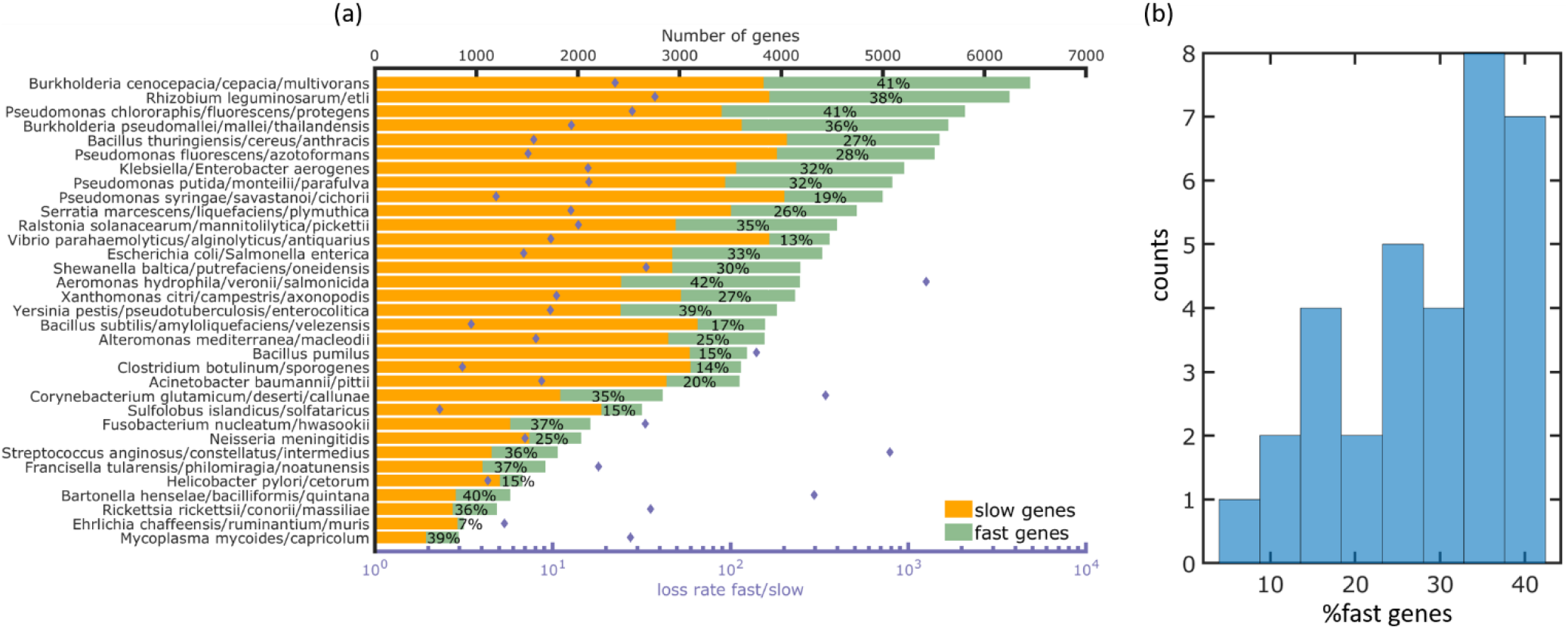
Inferred model parameters for the 33 prokaryotic clusters, as obtained by fitting the IGP-CGS model to mean genome intersections. Fitted model parameters are listed in Table S1. **a)** Top x-axis: the number of fast evolving and slow evolving genes in each cluster of genomes. The percentage of fast evolving genes is indicated for each cluster. Bottom x-axis: the ratio between fast and slow gene turnover rates. **b)** Histogram of the percentage of fast evolving genes across the 33 genomes clusters.

Fits of the model for the *E. coli* cluster are shown in Fig. 3. First, we fit the model directly to the gene commonality distribution, as it is usually done in the analyses of the U-shaped distributions [5-8]. Although a good fit to the U-shape distribution can be obtained (Fig. 3a), the inferred gene turnover rates are underestimated, as indicated by the poor fit obtained for the genome intersections (Fig. 3). Accordingly, the core genome size is overestimated by this fit. By contrast, when the turnover rates are inferred directly from gene intersections (Fig. 3b), the core genome size is estimated accurately but the number of singletons is overestimated (Fig. 3). This discrepancy is not unique to *E. coli* and was observed across the entire analyzed set of genome clusters (Fig.5). Specifically, under the IGP-CGS model, for all genome clusters, the goodness of fit of the model to mean intersections 〈*I*〉_*k*_ was greater than 0.995 (Fig. 5a), and the error in the core genome size was negligible (Fig. 5b), but the number of singletons was consistently overestimated (Fig. 5c).

**Figure 5.**
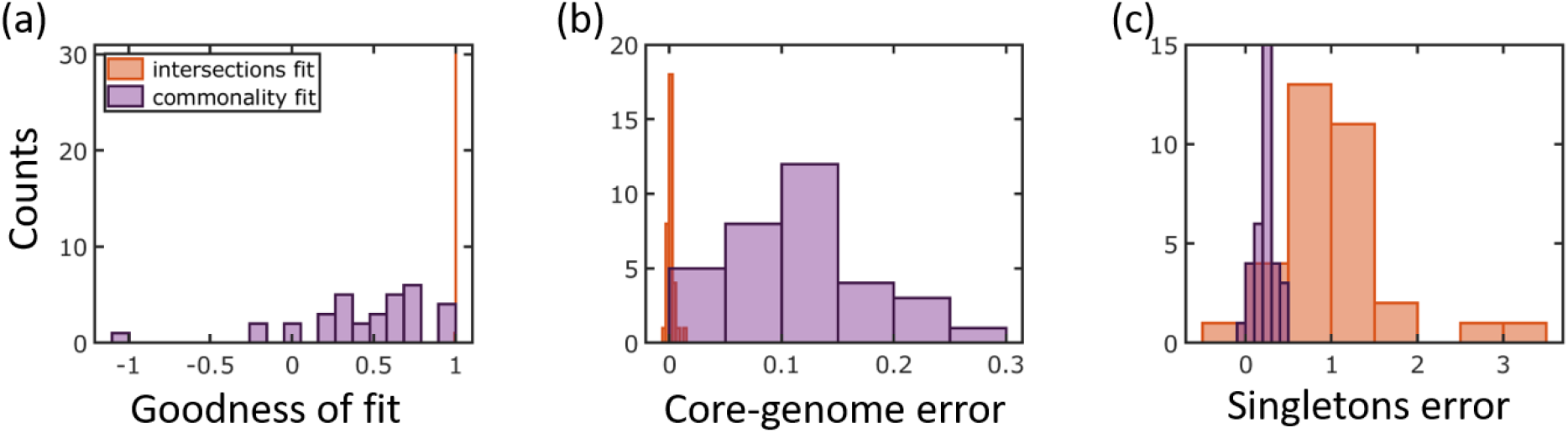
Comparison of the statistics of the IGP-CGS model fit to the 33 genomic clusters. a) Histogram for the goodness of fit *R^2^* to empirical mean genomes intersections *〈I〉_k_*. b) Histogram for the error in core-genome sizes *g_N_* of model fit. The error is calculated as 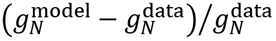. c) Histogram of the error in model prediction for the number of singletons, as computed from model mean genomes intersections using Eq. 1. The error is calculated as in panel **b**.

The fact that the model overestimates the number of singletons implies that fewer new genes are gained by evolving prokaryotic genomes than expected from the number of gene losses inferred directly from genomes intersections. This conclusion is supported by the direct fit of the model to the gene commonality distribution (Fig. 3): in this case, gene loss rates are underestimated, and as a result, the error shifts from the number of singletons to the core genome size (Fig. 5).

The exponential decay of genome intersections was obtained under the IGP-CGS assumptions whereby, on average, any lost gene is replaced by another gene and any gained gene is new to the given set of genomes. The overestimation of the number of singletons implies breakdown of either of these assumptions, or both. In other words, either the genome size is not constant, so that genomes shrink with time, or gene regain from the external gene pool is common (the gene pool is, then, finite), or both. Incorporating either a finite gene pool or a varying genome size into the calculation of genome intersections dramatically complicates the calculations and is beyond the scope of this work. Instead, we used simulations to explore separately the effect of genome size variation (IGP-V(ariable)GS assumption) and finite gene pool (F(inite)GP-CGS), and demonstrate for *E. coli* that a good fit to the genomic data can be achieved with either of these modifications.

To simulate the evolution of the *E. coli* pangenome for a finite gene pool (FGP-CGS) or a varying genome size (IGP-VGS), we used the same simulation scheme as before (Figs. 2e-g). Finite gene pool is introduced by limiting the number of genes that are available for the evolving genomes (see Methods). The violations of either of the IGP-CGS assumptions are considered separately to reduce the dimensionality of the parameter space and thus to allow realistic computation times. Given that it is impractical to scan numerically the entire 6-dimensional parameter spaces of the FGP-CGS or IGP-VGS models, the model parameters that were inferred for *E. coli* under the IGP-CGS assumptions were taken as the starting point (Figs. 2e-g, Table S1), and only a two-dimensional parameter space was scanned. For the FGP-CGS simulation, the two parameters were the sizes of the external gene pools, those for the slow and fast evolving genes. For the IGP-VGS simulation, the two parameters were the gain rate to loss rate ratios for the slow and fast evolving genes. Two examples for simulated datasets with genome intersection and gene commonality values similar to those in *E. coli* are shown in Fig. 6. Fitting the IGP-CGS model to the simulated datasets further demonstrated similarities between the simulated and genomic data. As with the genomic dataset, fitting the model directly to the genes commonality distribution resulted in underestimation of the gene turnover rates and the ensuing overestimation of the core genome size (Figs. 5a and 5c). Conversely, direct inference of turnover rates from the genome intersections resulted in the overestimation of the number of singletons (Figs. 6b and 6d). The discrepancy in the IGP-CGS model predictions, depending on the method of gene turnover rate inference, that was observed both with the simulated and the genomic datasets, indicates that violation of either of the model assumption is a plausible explanation for the overestimation of either, the number of singletons or the core genome size, by the IGP-CGS model.

**Figure 6.**
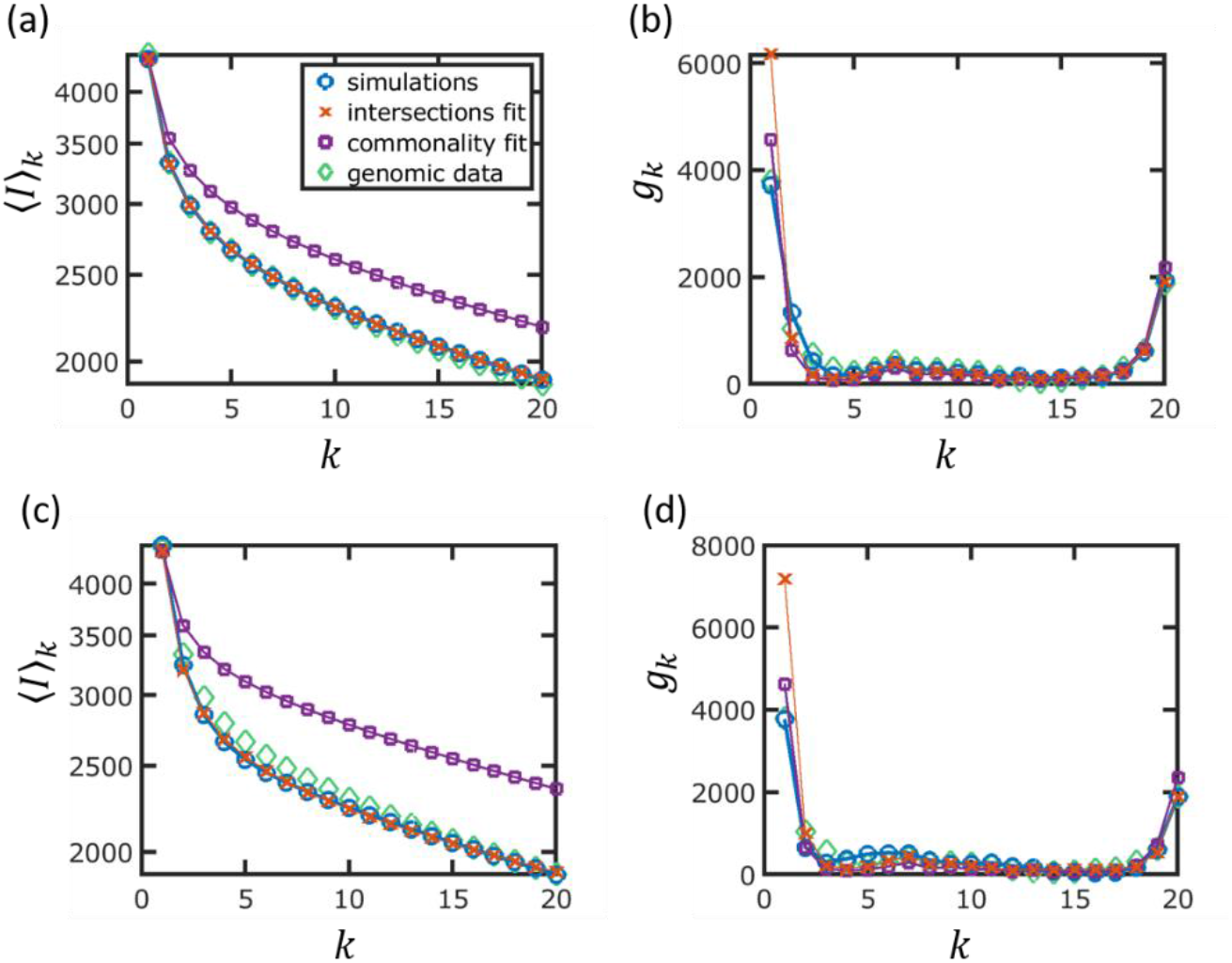
Analysis of simulated datasets for a finite gene pool (panels **a-b**) and a varying genome size (panels **c-d**). a) Mean genomes intersections for finite gene pool simulation. The parameters that were used in the simulations are the model parameters inferred under the IGP-CGS assumptions (see Table S1). For slow evolving genes the pool size was 3 times the number of slow genes, and the fast evolving gene pool was 9 times the number of fast genes. The IGP-CGS model fits and genomic data are also shown, as indicated in the legend. **b**) Gene commonality distribution for the finite gene pool simulation. **c**) Mean genome intersections for varying genome size. For slow evolving genes the ratio of gain and loss rates is 1.26, and for fast evolving genes that ratio of gain and loss rates is 0.16.

## Discussion

Here we present a theoretical framework to analyze quantitatively the evolutionary dynamics of microbial pangenomes and gene frequencies in them. Specifically, we infer genes turnover rates from genome intersections and reconstruct the U-shaped distribution that is observed for genes commonality for a genomic dataset consisting of 33 groups of closely related prokaryotic genomes. The asymmetrical U shape of the gene commonality distribution is extremely general, effectively, a universal of genome evolution, being observed for genomes at all phyletic depths, from a single species to the entirety of bacteria and archaea [4] [7]. By contrast, the pangenome size is not a well-defined characteristic, being sensitive to the number of genomes in a cluster, genome sampling and the tree depth [27]. It is therefore unclear what can be considered a ‘large’ or a ‘small’ pangenome, and comparison of the pangenome sizes for different organisms is not a straightforward task. Our analysis implies that, to attain insights into the evolutionary dynamics of microbes, it is preferable to measure and compare genes turnover rates that are more robust with respect to the number and sampling of the analyzed genomes.

Although our model produces good fits to genome intersection and the gene commonality distributions under the IGP-CGS assumptions, the number of singletons is systematically overestimated. It should be stressed that this overestimation cannot be explained by mis-annotations of genes, e.g., either under-prediction or over-prediction of poorly conserved, small genes that comprise the majority of the singletons. Prediction of more (or fewer) singletons will increase the genome size, which will affect the entire calculation and will not improve the fit. Thus, we hypothesized that the number of singletons is overestimated due to the violation of one or both of the IGP-CGS, namely, the infinite external gene pool and/or the (statistical) equality of the gene gain and loss rate, that is, the equilibrium state of the evolving microbial genomes. To test this hypothesis, we generated and analyzed simulated datasets for either finite gene pool (FGP-CGS) or varying genome sizes (IGP-VGS), obtaining results similar to those obtained in the genomic data analysis. However, numerical scanning of the two-dimensional parameter space for both FGP-CGS and IGP-VGS models failed to identify a single set of parameters yielding the best fit (Fig. S7). Furthermore, to make the computation feasible, violations of the IPG-CGS assumptions were introduced one at a time. Thus, the present analyses do not allow us to differentiate between the violations of the infinite gene pool and constant genome size assumptions. Furthermore, it appears likely that different factors are important in different groups of microbes and that, in some cases, both assumptions are violated simultaneously. Indeed, there is empirical evidence of deviations from each of these assumptions in bacterial evolution. Our previous analysis performed with the same ATGC dataset has shown that the evolution of most groups of bacteria was dominated by genes loss, the rate of which far exceeded that of gene gain [13]. Conceivably, some of these groups are headed for extinction whereas in others, the steady gene loss is compensated by episodes of massive gene gain. Thus, the constant genome size assumption appears to be frequently violated. Furthermore, direct measurements of gene regain have shown conspicuous differences among prokaryotes, such that, for some bacteria, the pool available for gene acquisition appeared to be effectively infinite, whereas for others, it was found to be finite, and comparatively small [13]. Accordingly, the infinite gene pool assumption is, in the least, not universally valid either.

The breakdown of both assumptions of IGP-CGS reflects the cost of incorporation of new genes by microbes. In both cases, the number of gained new genes (as opposed to variants of genes already represented in a genome) is smaller than that predicted by the IGP-CGS model, suggesting that the major factor that restricts the expansion of the pangenome is the capacity to successfully incorporate new genes, i.e. the HGT barrier [26]. The existence of a substantial HGT barrier is supported also by experimental analysis of the assimilation of xenologs of orthologous genes by bacteria [28]. In these experiments, replacement of an essential *E. coli* gene by xenologs resulted in a substantial drop in fitness that was partially alleviated during subsequent evolution of the recipient bacteria in the laboratory. In the course of long term evolution of microbes, the HGT barrier is likely to be broken episodically as a result of major changes in environmental conditions when extensive HGT favors survival [13, 29]. Different functional classes of microbial genes show substantial differences in evolutionary plasticity, or in other words, are differentially affected by the HGT barrier [20]. Thus, analysis of pangenome dynamics separately for each class can be expected to reveal the interplay between the key factors of genome evolution.

## Methods

### Model for genomes intersections evolution

The evolution of the genome size can be modelled as a random process, where genes are gained and lost stochastically, with rates *P*^+^ and *P*−, respectively. Accordingly, the dynamics of the number of genes *x* is given by the equation [24]

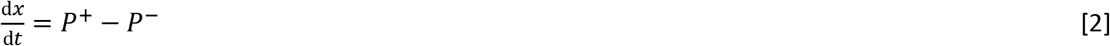

For IGP-CGS model, genomes intersection of *k* genomes is given by [19]

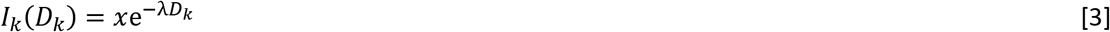

where the decay constant is given by the per-gene loss rate *λ* = *P*^−^/*x*, and *D_k_* is the total evolutionary distance spanned by those *k* genomes. Specifically, *D_k_* is given by the sum of all branch lengths in the phylogenetic tree that describes the evolutionary relations of the *k* genomes. Analyses of the genomic data imply that genomes are composed of slow and fast evolving genes [21]. Thus,

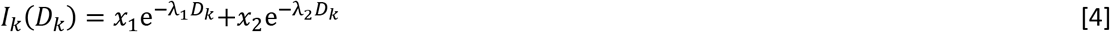

with *x*_1_ and *x*_2_ being the average numbers of genes in each class. Fitting the data to the model requires therefore inference of four parameters: *x*_1_ *x*_2_, *λ*_1_ and *λ*_2_.

For a cluster of *N* genomes there are 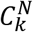 subsets of *k* genomes, where 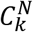 denotes the binomial coefficient. To extract genes commonality from genomes intersections using Eq. 1, we wish to evaluate the mean intersection for each *k*, denoted 〈*I*〉_*k*_. The first stage is to obtain all possible 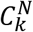 phylogenetic subtrees that include *k* genomes, by pruning the tree of the complete set of *N* genomes. Next, the sum of the branch lengths for each *k* genomes subtree is evaluated to construct the probability density function *P*(*D_k_*) to observe the sum of branch lengths between *D_k_* and *D_k_* + *dD_k_*. Finally, the mean intersection 〈*I*〉_*k*_ is given by a weighted average over branch lengths

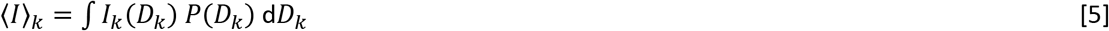

where *I_k_*(*D_k_*) is given by either Eq. 3 or Eq. 4. Note that *P*(*D_k_*) reflects both, the phylogenetic tree topology and the branch lengths.

### Simulation scheme

A cluster of genomes is simulated along a tree, using the Gillespie simulations scheme [30]. A genome is represented as a collection of genes, and possible mutations are either gene gain or gene loss of rates *P*^+^ and *P*^−^, respectively. The occurrence of mutations therefore follows a Poisson distribution with parameter *a*_0_, which is given by

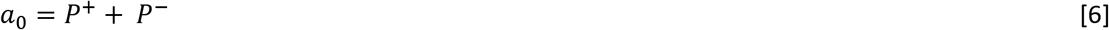

The waiting time between mutation events *τ* follows an exponential distribution with rate parameter *a*_0_. At each simulation step *τ* is picked from an exponential distribution, using

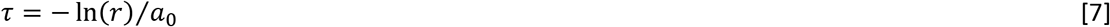

where *r* is a random number drawn from a uniform distribution between 0 and 1. After determining the waiting time, it is determined whether a gene is gained or lost at random, according to *P*^+^ and *P*^−^. Under the IGP-CGS assumptions the simulations scheme takes as an input in addition to the tree two parameters: initial genome size *x*, and gene loss rate *P*^−^ (which in under the CGS assumption also determines the gain rate *P*^+^).

Empirical genome intersections imply that two gene turnover rates are required to fit the genomic data (see Eq. 4). To generate more realistic datasets, genomes containing two classes of genes, fast and slow evolving genes, are simulated. In addition to the tree, this simulation scheme includes four parameters: *x*_1_/*x*_2_ and loss rates for slow and fast evolving genes, 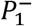 and 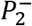. Extending the simulation scheme to account for breakdown of either of the IGP-CGS assumptions will add two parameters, to a total of six. For the FGP-CGS assumption, the total number of genes that are available to the evolving genomes is set to a finite number. Fast and slow genes evolve independently and are assumed to be drawn from two different pools, such that the two additional parameters are the gene pool sizes, *L*_1_ and *L*_2_. Under the IGP-VGS assumption, the gain and loss rates are not necessarily equal, which add two additional parameters, 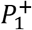 and 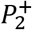.

### Genomic dataset

We used the Alignable Tight Genomic Clusters (ATGC) database [31] to compile 33 groups of closely related prokaryotic genome. The dataset includes 32 groups (or ATGCs) of bacteria and one group of archaea (see Table S1 for the list of ATGCs analyzed in this study). The selected ATGCs meet the following criteria: i) maximum pairwise tree distance is at least 0.1 substitutions per site, and ii) the phylogenetic tree contains more than two clades, such that pairwise tree distances are centered around more than two typical values. To allow reasonable computational times, ATGCs with more than 20 genomes were sampled such that each ATGC contains at most 20 representative genomes. Throughout the analysis, phylogenetic trees were rescaled to compensate for the systematic underestimation of branch lengths that is apparently due to homologous recombination [19].

## Acknowledgements

We thank Koonin group members for helpful discussions. The authors’ research is supported by intramural research program funds of the National Institutes of Health (National Library of Medicine). This work utilized the computational resources of the NIH HPC Biowulf cluster (http://hpc.nih.gov).

## References

[1] G. Vernikos, D. Medini, D.R. Riley, H. Tettelin, Ten years of pan-genome analyses, Curr Opin Microbiol, 23 (2015) 148–154.

[2] J.O. McInerney, A. McNally, M.J. O’Connell, Why prokaryotes have pangenomes, Nat Microbiol, 2 (2017) 17040.

[3] D. Medini, C. Donati, H. Tettelin, V. Masignani, R. Rappuoli, The microbial pan-genome, Curr Opin Genet Dev, 15 (2005) 589–594.

[4] Y.I. Wolf, K.S. Makarova, N. Yutin, E.V. Koonin, Updated clusters of orthologous genes for Archaea: a complex ancestor of the Archaea and the byways of horizontal gene transfer, Biol Direct, 7 (2012) 46.

[5] F. Baumdicker, W.R. Hess, P. Pfaffelhuber, The infinitely many genes model for the distributed genome of bacteria, Genome Biol Evol, 4 (2012) 443–456.

[6] R.E. Collins, P.G. Higgs, Testing the infinitely many genes model for the evolution of the bacterial core genome and pangenome, Mol Biol Evol, 29 (2012) 3413–3425.

[7] A.E. Lobkovsky, Y.I. Wolf, E.V. Koonin, Gene frequency distributions reject a neutral model of genome evolution, Genome Biol Evol, 5 (2013) 233–242.

[8] A. Mazzolini, M. Gherardi, M. Caselle, M. Cosentino Lagomarsino, M. Osella, Statistics of Shared Components in Complex Component Systems, Physical Review X, 8 (2018).

[9] B. Haegeman, J.S. Weitz, A neutral theory of genome evolution and the frequency distribution of genes, BMC Genomics, 13 (2012) 196.

[10] E.V. Koonin, K.S. Makarova, L. Aravind, Horizontal gene transfer in prokaryotes: quantification and classification, Annu Rev Microbiol, 55 (2001) 709–742.

[11] C. Pal, B. Papp, M.J. Lercher, Adaptive evolution of bacterial metabolic networks by horizontal gene transfer, Nat Genet, 37 (2005) 1372–1375.

[12] T.J. Treangen, E.P. Rocha, Horizontal transfer, not duplication, drives the expansion of protein families in prokaryotes, PLoS Genet, 7 (2011) e1001284.

[13] P. Puigbo, A.E. Lobkovsky, D.M. Kristensen, Y.I. Wolf, E.V. Koonin, Genomes in turmoil: quantification of genome dynamics in prokaryote supergenomes, BMC Biol, 12 (2014) 66.

[14] W.F. Doolittle, Lateral genomics, Trends Cell Biol, 9 (1999) M5–8.

[15] M.A. Brockhurst, E. Harrison, J.P.J. Hall, T. Richards, A. McNally, C. MacLean, The Ecology and Evolution of Pangenomes, Current Biology, 29 (2019) R1094–R1103.

[16] M.R. Domingo-Sananes, J.O. McInerney, Selection-based model of prokaryote pangenomes, bioRxiv, DOI 10.1101/782573(2019).

[17] M. Kimura, J.F. Crow, The Number of Alleles That Can Be Maintained in a Finite Population, Genetics, 49 (1964) 725–738.

[18] M.A. Moldovan, M.S. Gelfand, Pangenomic Definition of Prokaryotic Species and the Phylogenetic Structure of Prochlorococcus spp, Front Microbiol, 9 (2018) 428.

[19] J. Iranzo, Y.I. Wolf, E.V. Koonin, I. Sela, Gene gain and loss push prokaryotes beyond the homologous recombination barrier and accelerate genome sequence divergence, Nat Commun, 10 (2019) 5376.

[20] I. Sela, Y.I. Wolf, E.V. Koonin, Selection and Genome Plasticity as the Key Factors in the Evolution of Bacteria, Physical Review X, 9 (2019).

[21] Y.I. Wolf, K.S. Makarova, A.E. Lobkovsky, E.V. Koonin, Two fundamentally different classes of microbial genes, Nat Microbiol, 2 (2016) 16208.

[22] G. Plata, C.S. Henry, D. Vitkup, Long-term phenotypic evolution of bacteria, Nature, 517 (2015) 369–372.

[23] T.N. Mavrich, G.F. Hatfull, Bacteriophage evolution differs by host, lifestyle and genome, Nat Microbiol, 2 (2017) 17112.

[24] I. Sela, Y.I. Wolf, E.V. Koonin, Theory of prokaryotic genome evolution, Proc Natl Acad Sci U S A, 113 (2016) 11399–11407.

[25] L. Comtet, Advanced Combinatorics, 1974.

[26] R. Sorek, Y. Zhu, C.J. Creevey, M.P. Francino, P. Bork, E.M. Rubin, Genome-Wide Experimental Determination of Barriers to Horizontal Gene Transfer, Science, 318 (2007) 1449–1452.

[27] H. Tettelin, V. Masignani, M.J. Cieslewicz, C. Donati, D. Medini, N.L. Ward, S.V. Angiuoli, J. Crabtree, A.L. Jones, A.S. Durkin, R.T. DeBoy, T.M. Davidsen, M. Mora, M. Scarselli, I. Margarit y Ros, J.D. Peterson, C.R. Hauser, J.P. Sundaram, W.C. Nelson, R. Madupu, L.M. Brinkac, R.J. Dodson, M.J. Rosovitz, S.A. Sullivan, S.C. Daugherty, D.H. Haft, J. Selengut, M.L. Gwinn, L. Zhou, N. Zafar, H. Khouri, D. Radune, G. Dimitrov, K. Watkins, K.J.B. O’Connor, S. Smith, T.R. Utterback, O. White, C.E. Rubens, G. Grandi, L.C. Madoff, D.L. Kasper, J.L. Telford, M.R. Wessels, R. Rappuoli, C.M. Fraser, Genome analysis of multiple pathogenic isolates of Streptococcus agalactiae: Implications for the microbial “pan-genome”, Proceedings of the National Academy of Sciences, 102 (2005) 13950–13955.

[28] S. Bershtein, A.W. Serohijos, S. Bhattacharyya, M. Manhart, J.M. Choi, W. Mu, J. Zhou, E.I. Shakhnovich, Protein Homeostasis Imposes a Barrier on Functional Integration of Horizontally Transferred Genes in Bacteria, PLoS Genet, 11 (2015) e1005612.

[29] Y.I. Wolf, E.V. Koonin, Genome reduction as the dominant mode of evolution, BioEssays, 35 (2013) 829–837.

[30] D.T. Gillespie, Exact stochastic simulation of coupled chemical reactions, The Journal of Physical Chemistry, 81 (1977) 2340–2361.

[31] D.M. Kristensen, Y.I. Wolf, E.V. Koonin, ATGC database and ATGC-COGs: an updated resource for micro- and macro-evolutionary studies of prokaryotic genomes and protein family annotation, Nucleic Acids Res, 45 (2017) D210–D218.

