## Supplementary Information for "Horizontal gene transfer barrier shapes the evolution of prokaryotic pangenomes"

**Contents**

**I. Supplementary Figures** 2

**II. Supplementary Tables** 9

### I. SUPPLEMENTARY FIGURES

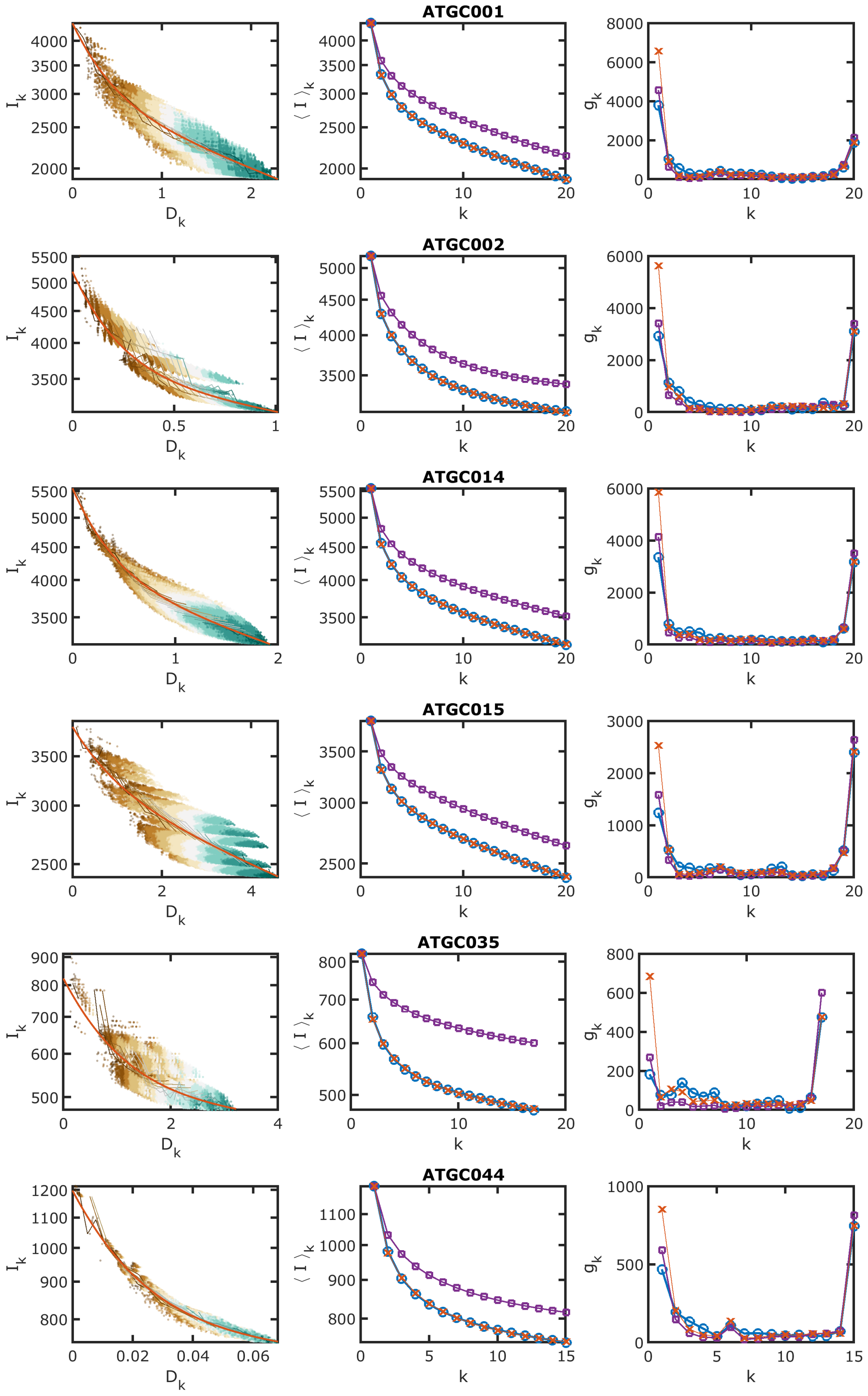


FIG. S1: Genomes intersections and genes commonality distribution for the analyzed genomic dataset. The IGP-CGS model fits are also indicated, see legend of Fig. 1a in the main text. Each row shows different genomes cluster, the ATGC number is indicated.

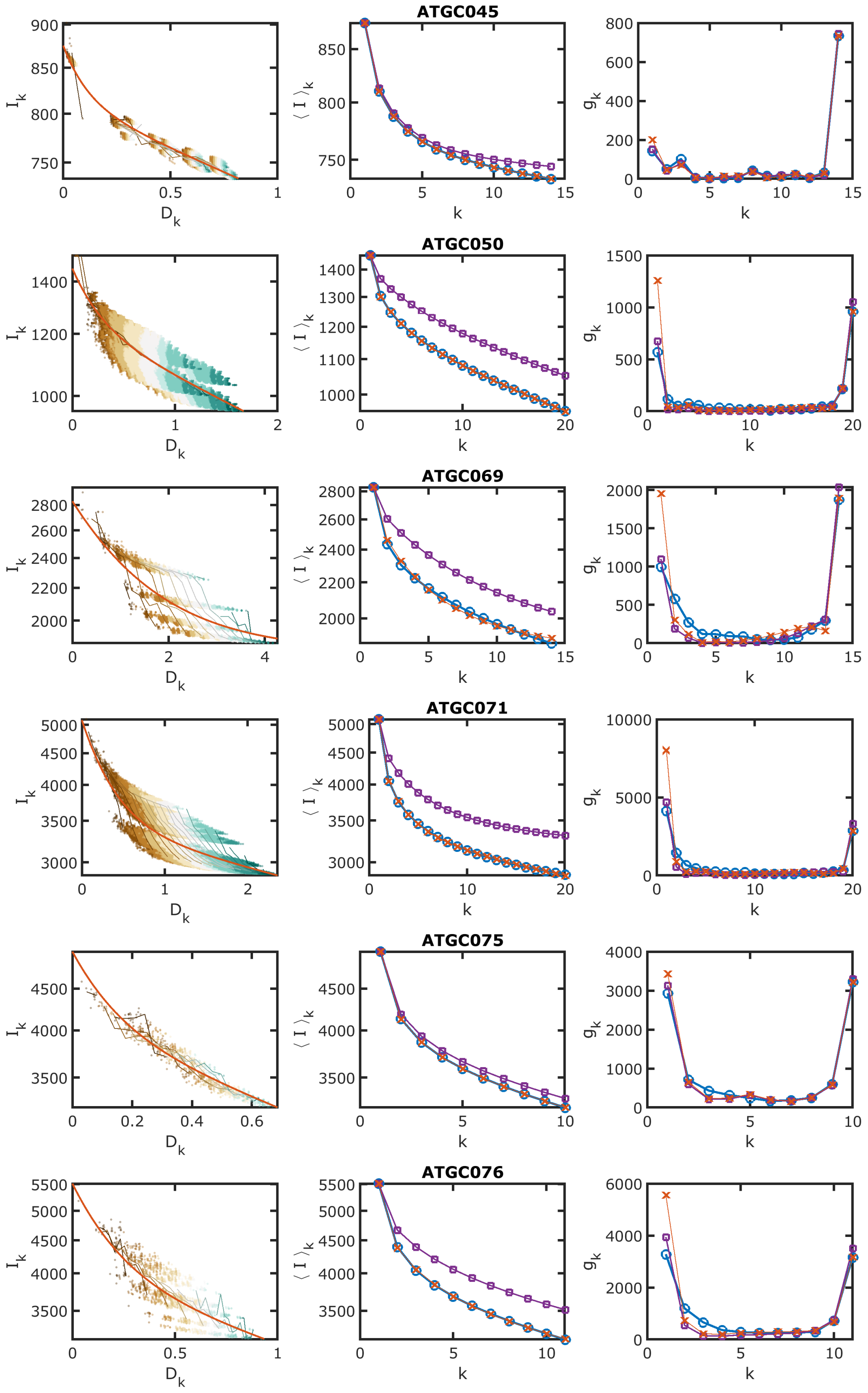


FIG. S2: Same as Fig. S1.

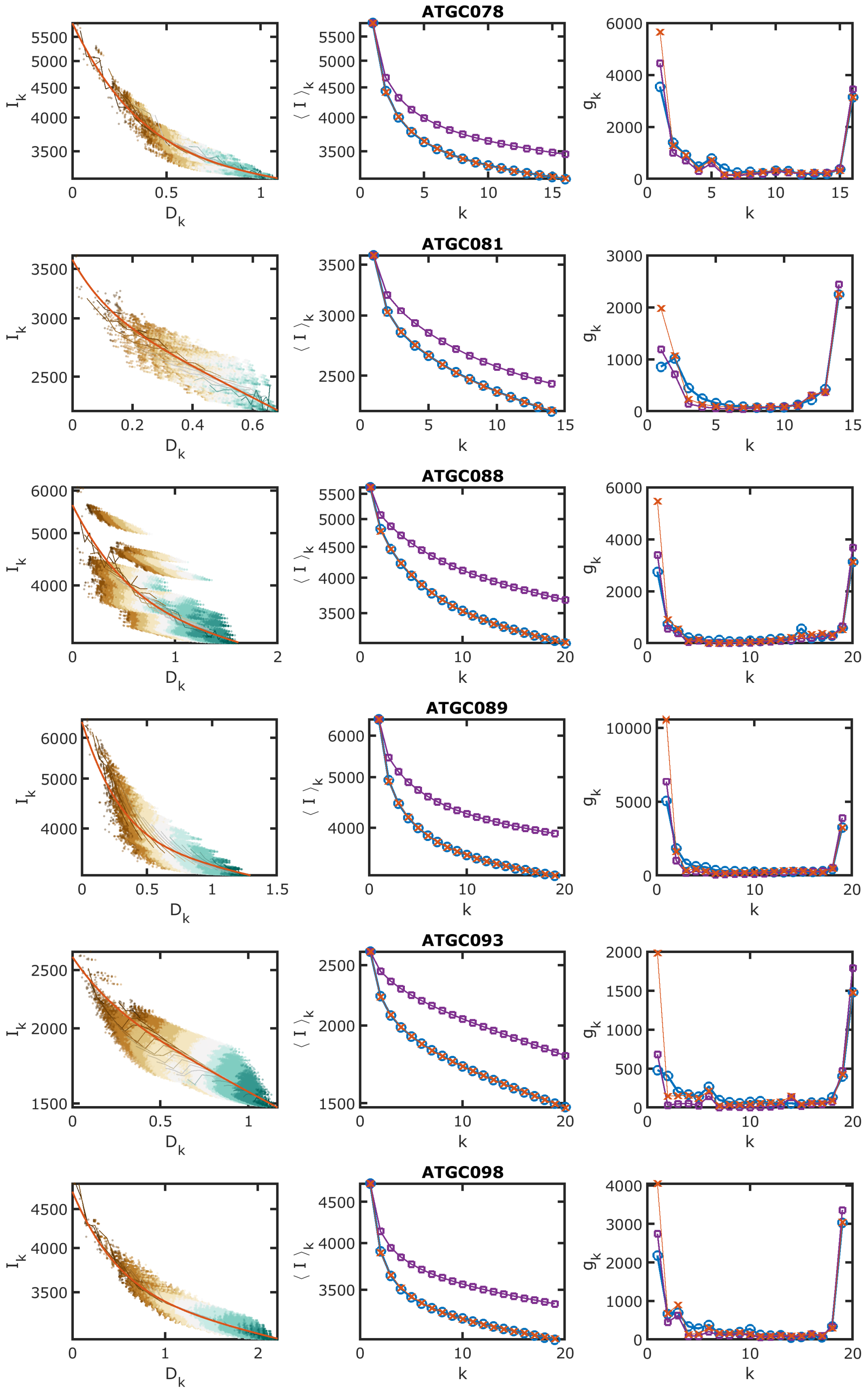


FIG. S3: Same as Fig. S1.

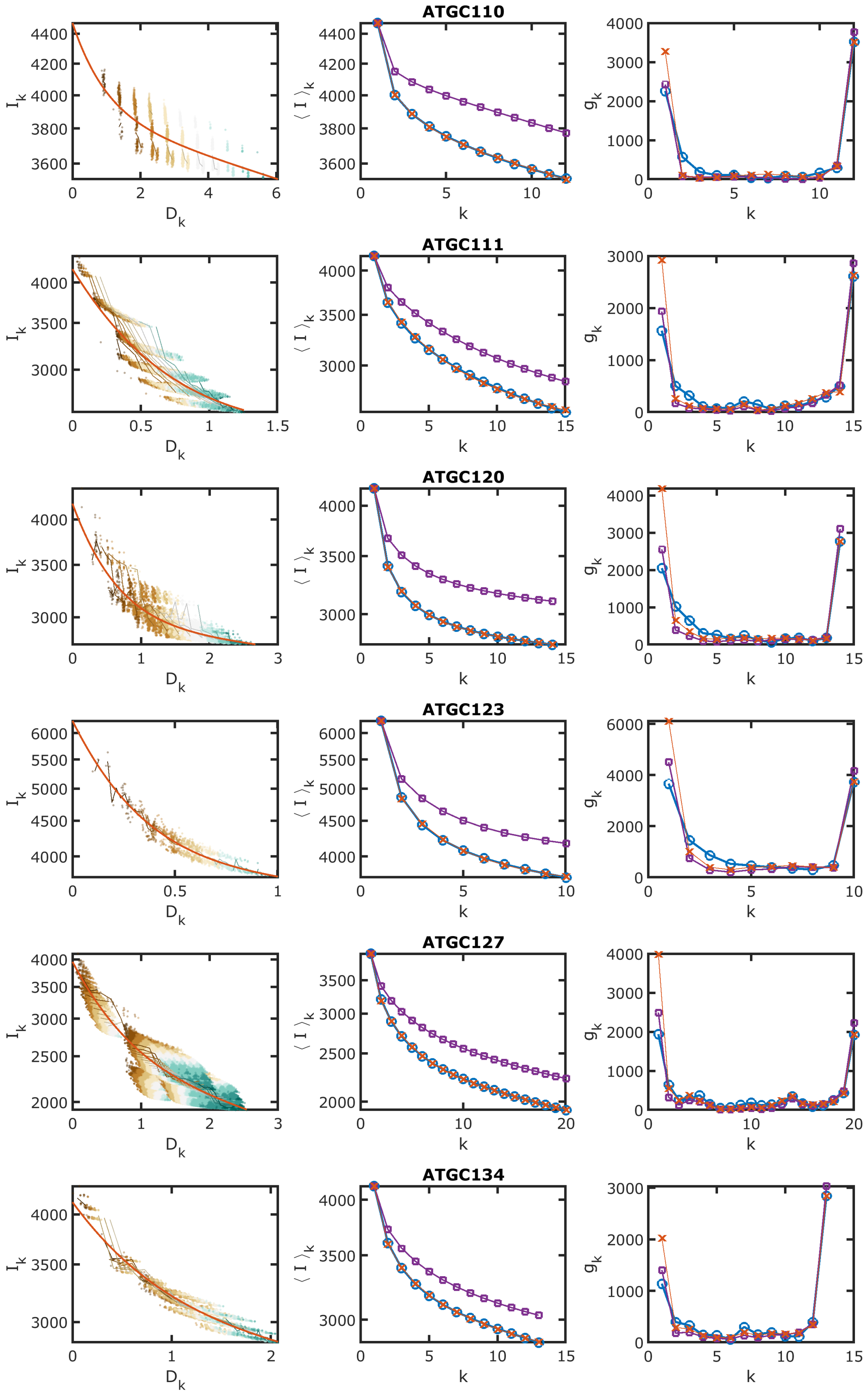


FIG. S4: Same as Fig. S1.

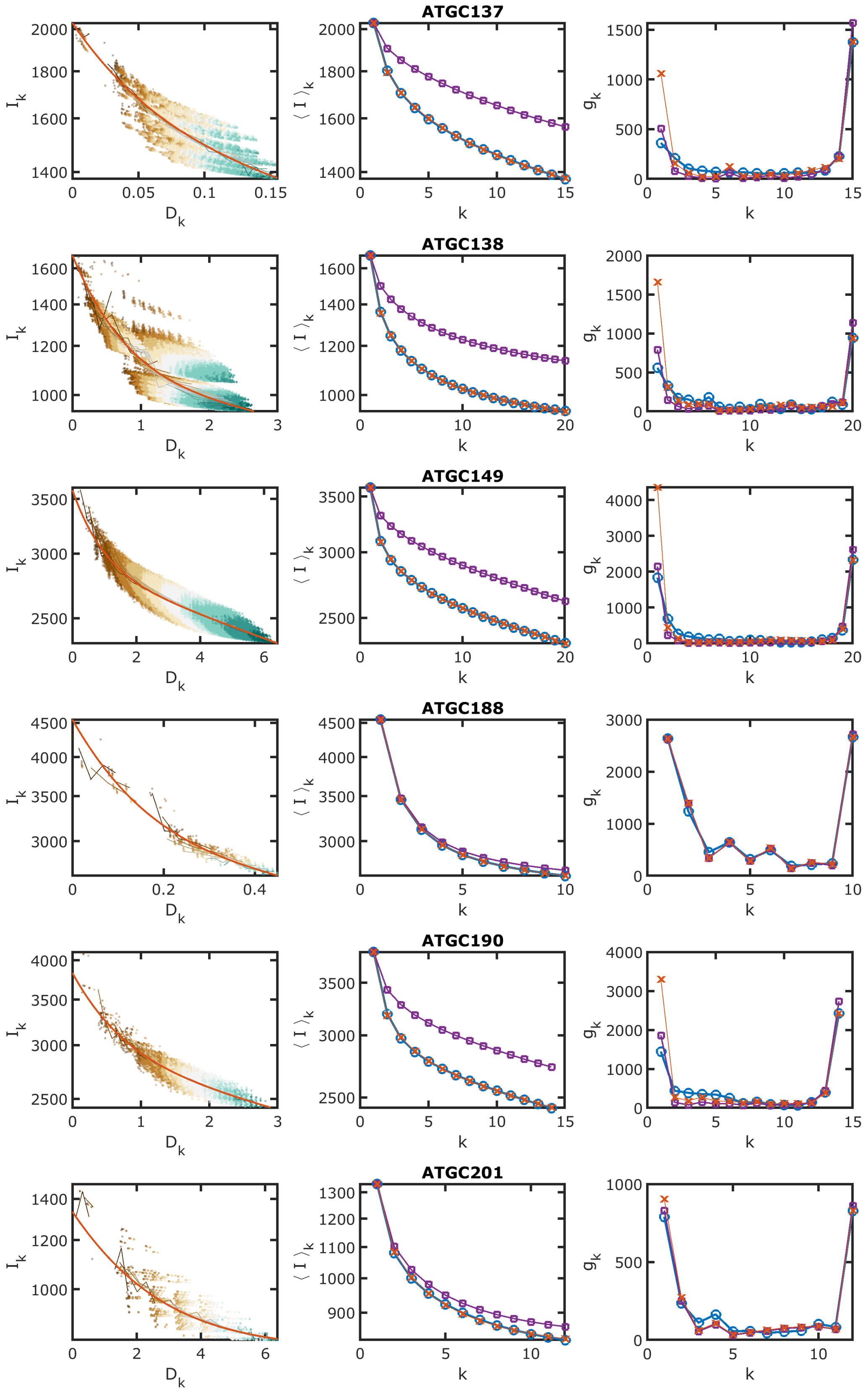


FIG. S5: Same as Fig. S1.

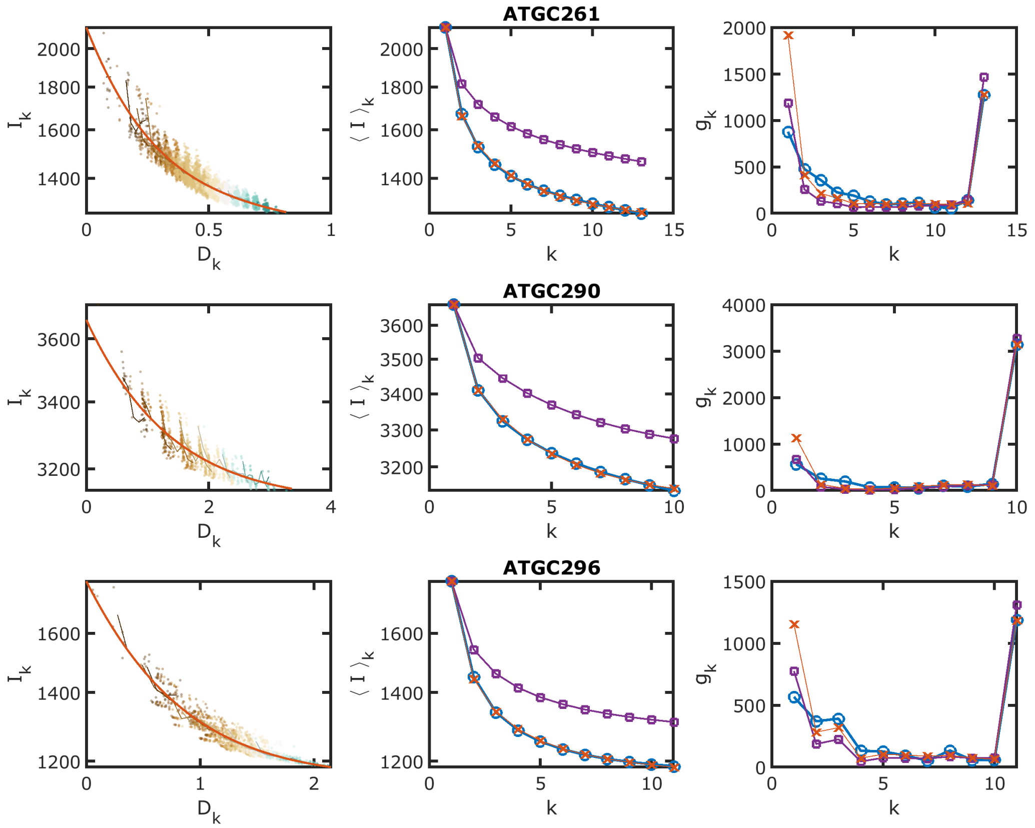


FIG. S6: Same as Fig. S1.

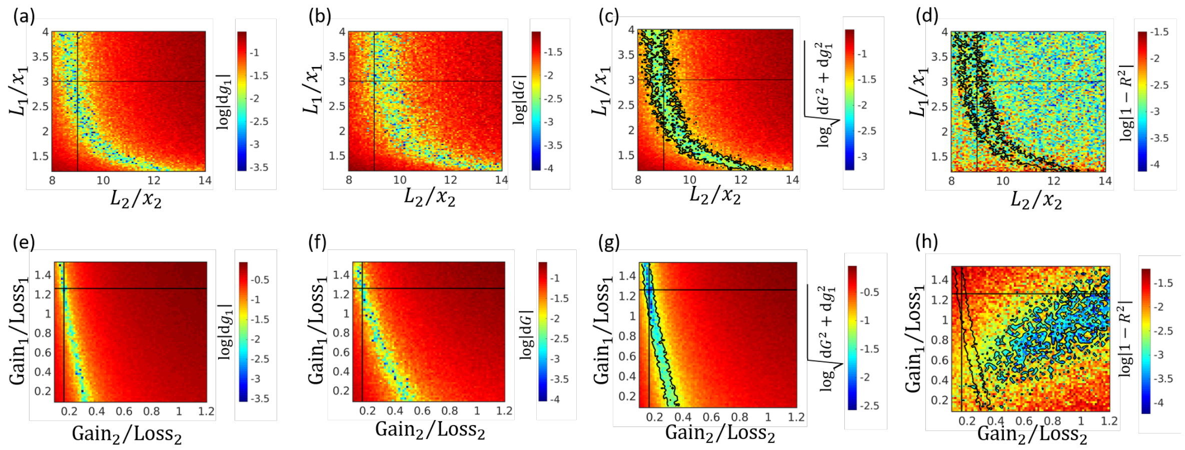


FIG. S7: The similarity of simulated datasets to the genomic data of ATGC001. Simulations for different pool sizes under the FGP-CGS assumption are shown in panels a-d. Simulations for different gain to loss rations under the IGP-VGS assumption are shown in panels e-h. The similarity between the simulated data is quantified by the error in the number of singletons (panels a and e), the error in the pangenome size (panels b and f), a combined measure of the number of singletons and the pangenome size (panels c and g), and the goodness of fit for the mean intersections (panels d and h). the error $dX$ is calculated as $(X_{model} - X_{data}/X_{data}$. Contour lines of the optimal combined measure are indicated in panels c and g. For comparison, the optimal region in terms of the combined measure is also shown in panels d and h. The parameters that were used in the simulations that are shown in Fig. 5 of the main text are indicated by a vertical and a horizontal black lines in all panels.

| Genomes names | ATGC | # genomes | # slow genes | # fast genes | $P^-_{slow}$ | $P^-_{fast}$ |
|---|---|---|---|---|---|---|
| Escherichia coli/Salmonella enterica | ATGC001 | 20 | 2929 | 1471 | 563 | 3877 |
| Klebsiella/Enterobacter aerogenes | ATGC002 | 20 | 3558 | 1648 | 502 | 7962 |
| Bacillus thuringiensis/cereus/anthracis | ATGC014 | 20 | 4057 | 1497 | 519 | 4081 |
| Bacillus subtilis/amyloliquefaciens/velezensis | ATGC015 | 20 | 3178 | 658 | 195 | 679 |
| Mycoplasma mycoides/capricolum | ATGC035 | 17 | 504 | 319 | 12 | 341 |
| Rickettsia rickettsii/conorii/massiliae | ATGC044 | 15 | 768 | 429 | 581 | 20706 |
| Ehrlichia chaffeensis/ruminantium/muris | ATGC045 | 14 | 814 | 62 | 102 | 550 |
| Helicobacter pylori/cetorum | ATGC050 | 20 | 1232 | 222 | 187 | 812 |
| Corynebacterium glutamicum/deserti/callunae | ATGC069 | 14 | 1828 | 1002 | 2 | 625 |
| Pseudomonas putida/monteilii/parafulva | ATGC071 | 20 | 3452 | 1636 | 278 | 4457 |
| Pseudomonas syringae/savastanoi/cichorii | ATGC075 | 10 | 4031 | 965 | 1343 | 6464 |
| Pseudomonas fluorescens/azotoformans | ATGC076 | 11 | 3958 | 1552 | 975 | 7120 |
| Pseudomonas chlororaphis/fluorescens/protegens | ATGC078 | 16 | 3418 | 2388 | 308 | 8651 |
| Clostridium botulinum/sporogenes | ATGC081 | 14 | 3112 | 488 | 1477 | 4604 |
| Burkholderia pseudomallei/mallei/thailandensis | ATGC088 | 20 | 3612 | 2028 | 357 | 4552 |
| Burkholderia cenocepacia/cepacia/multivorans | ATGC089 | 19 | 3829 | 2618 | 501 | 11276 |
| Sulfolobus islandicus/solfataricus | ATGC093 | 20 | 2230 | 397 | 792 | 1835 |
| Serratia marcescens/liquefaciens/plymuthica | ATGC098 | 19 | 3505 | 1235 | 228 | 2895 |
| Vibrio parahaemolyticus/alginolyticus/antiquarius | ATGC110 | 12 | 3886 | 587 | 65 | 637 |
| Aeromonas hydrophila/veronii/salmonicida | ATGC111 | 15 | 2426 | 1757 | 2 | 3062 |
| Shewanella baltica/putrefaciens/oneidensis | ATGC120 | 14 | 2929 | 1259 | 66 | 2217 |
| Rhizobium leguminosarum/etli | ATGC123 | 10 | 3883 | 2362 | 217 | 8186 |
| Yersinia pestis/pseudotuberculosis/enterocolitica | ATGC127 | 20 | 2420 | 1535 | 234 | 2270 |
| Xanthomonas citri/campestris/axonopodis | ATGC134 | 13 | 3014 | 1119 | 130 | 1368 |
| Neisseria meningitidis | ATGC137 | 15 | 1521 | 510 | 1230 | 8592 |
| Francisella tularensis/philomiragia/noatunensis | ATGC138 | 20 | 1062 | 617 | 53 | 954 |
| Acinetobacter baumannii/pittii | ATGC149 | 20 | 2874 | 715 | 94 | 818 |
| Ralstonia solanacearum/mannitolilytica/pickettii | ATGC188 | 10 | 2963 | 1587 | 837 | 11668 |
| Alteromonas mediterranea/macleodii | ATGC190 | 14 | 2889 | 943 | 179 | 1441 |
| Bartonella henselae/bacilliformis/quintana | ATGC201 | 12 | 799 | 535 | 1 | 236 |
| Fusobacterium nucleatum/hwasookii | ATGC261 | 13 | 1338 | 782 | 106 | 3503 |
| Bacillus pumilus | ATGC290 | 10 | 3102 | 560 | 3 | 435 |
| Streptococcus anginosus/constellatus/intermedius | ATGC296 | 11 | 1152 | 647 | 1 | 908 |

TABLE S1: Genome clusters (ATGCs) in the analyzed dataset and model parameters that were inferred under the IGP-CGS assumptions.

### II. SUPPLEMENTARY TABLES
